# Opposing effects of acellular and whole cell pertussis vaccines on *Bordetella pertussis* biofilm formation, Siglec-F+ neutrophil recruitment and bacterial clearance in mouse nasal tissues

**DOI:** 10.1101/2024.01.23.576795

**Authors:** Jesse M. Hall, Jessica L. Gutiérrez-Ferman, Mohamed M. Shamseldin, Myra Guo, Yash A. Gupta, Rajendar Deora, Purnima Dubey

**Affiliations:** Department of Microbial Infection and Immunity, The Ohio State University, Columbus, OH; Department of Microbiology, The Ohio State University, Columbus, OH; Department of Microbiology and Immunology, Faculty of Pharmacy, Helwan University Ain Helwan, Helwan, 11795, Egypt

**Keywords:** *Bordetella pertussis*, whooping cough, mucosal immunology, vaccines, biofilms

## Abstract

Despite global vaccination, pertussis caused by *Bordetella pertussis* (*Bp*) is resurging. Pertussis resurgence is correlated with the switch from whole cell vaccines (wPV) that elicit T_H_1/T_H_17 polarized immune responses to acellular pertussis vaccines (aPV) that elicit primarily T_H_2 polarized immune responses. One explanation for the increased incidence in aPV-immunized individuals is the lack of bacterial clearance from the nose. To understand the host and bacterial mechanisms that contribute to *Bp* persistence, we evaluated bacterial localization and the immune response in the nasal associated tissues (NT) of naïve and immunized mice following *Bp* challenge. *Bp* resided in the NT of unimmunized and aPV-immunized mice as biofilms. In contrast, *Bp* biofilms were not observed in wPV-immunized mice. Following infection, Siglec-F+ neutrophils, critical for eliminating *Bp* from the nose, were recruited to the nose at higher levels in wPV immunized mice compared to aPV immunized mice. Consistent with this observation, the neutrophil chemokine CXCL1 was only detected in the NT of wPV immunized mice. Importantly, the bacteria and immune cells were primarily localized within the NT and were not recovered by nasal lavage (NL). Together, our data suggest that the T_H_2 polarized immune response generated by aPV vaccination facilitates persistence in the NT by impeding the infiltration of immune effectors and the eradication of biofilms In contrast, the T_H_1/T_H_17 immune phenotype generated by wPV, recruits Siglec-F+ neutrophils that rapidly eliminate the bacterial burden and prevent biofilm establishment. Thus, our work shows that aPV and wPV have opposing effects on *Bp* biofilm formation in the respiratory tract and provides a mechanistic explanation for the inability of aPV vaccination to control bacterial numbers in the nose and prevent transmission.

**Author Summary:** Acellular pertussis vaccine (aPV) immunized individuals maintain a nasal reservoir of *Bordetella pertussis* (*Bp*) and thus have the potential to transmit the infection to vulnerable individuals. Here we provide a mechanistic explanation for the inability of aPV to eliminate *Bp* from the nasal cavity. We show that following bacterial challenge of aPV immunized mice, Siglec-F+ neutrophils and other immune effectors are not recruited to the nose. Consequently, *Bp* remain in the nose and form biofilms. In contrast, whole cell pertussis (wPV) immunized mice produce immune effectors following bacterial challenge that recruit Siglec-F+ neutrophils to the nose. *Bp* burden is cleared from the nasal tissues, thereby preventing bacterial persistence and the formation of biofilms.

## Introduction

The respiratory disease known as whooping cough (pertussis) is one of the more prevalent vaccine preventable diseases, despite greater than 84% global vaccination rates (1). Pertussis is caused by infection of the respiratory epithelium by the Gram-negative bacterium *Bordetella pertussis* (*Bp*). Whole cell pertussis vaccines (wPV) significantly reduced the number of cases worldwide. However, due to their reactogenicity and resulting public health concerns, wPV were replaced in many countries by acellular pertussis vaccines (aPV) since the late 1990s (2, 3). Since the implementation of less reactogenic aPV, cyclic increases in the number of pertussis cases were observed (4). Animal studies showed that aPV prevent pertussis disease and clear the bacteria from the lungs, but do not prevent *Bp* infection and colonization of the nose (5–7). Thus, aPV-immunized individuals are asymptomatic carriers who serve as silent sources of *Bp* transmission (8–10). With ongoing use of aPV, the pool of asymptomatic carriers continues to increase.

We previously showed that *Bp* forms biofilms in the nose of naïve mice (11, 12). However, the mechanisms that permit *Bp* persistence in the nose following aPV immunization are undefined. To understand how *Bp* persists in the nose of aPV immunized mice and not in wPV immunized mice, we followed *Bp* colonization, and characterized the immune effectors elicited following bacterial challenge. *Bp* bacterial load was similar in unimmunized and aPV-immunized mice, and confocal microscopy showed that *Bp* formed biofilms in the nasal cavity of these mice. In contrast, *Bp* numbers were significantly reduced from the nose of wPV immunized mice and biofilms were not observed. Following *Bp* challenge, CD4+ T cells and Siglec-F+ neutrophils were recruited to the nose of wPV-immunized mice but were not found in the nose of unimmunized and aPV-immunized mice. Together, our data suggest that biofilm formation results in *Bp* persistence in aPV immunized animals and provide a mechanistic explanation for the development of a *Bp* nasal reservoir in immune individuals.

## Results

### aPV immunization fails to eradicate *Bp* biofilms in the mouse nose

Most animal studies have evaluated bacterial burden in the nose by washing the airway and collecting the nasal lavage (NL) (6, 7, 13, 14) while we enumerate bacterial load in the nasal tissues, which includes the septum, turbinates, and nasal epithelial cells (15). Here, we hypothesized that enumeration of bacterial burden in the NL alone vastly underrepresents the true *Bp* burden in the nose. To test this, we enumerated bacterial burden in both NL and in the nasal tissues (NT) in the same mice. We challenged C57BL/6 mice intranasally (IN) with the laboratory strain Bp536 (16). NL was collected by flushing the nares followed by dissection of the NT (**Fig. 1A**) which was then mechanically dissociated. The NT suspension and NL were diluted and plated to enumerate *Bp* CFUs. While bacteria were present in both compartments, bacterial load was 1 log higher in NT than NL on days 4 and 7, and 2 logs higher in NT on day 14 post-challenge (**Fig. 1 and Fig. S1A**). Most of the bacteria were present in the NT with approximately 6-12 percent of the total bacterial burden present in NL (**Fig. S1B**). Thus, *Bp* is primarily localized in the NT, not in the NL.

**Fig. 1.**
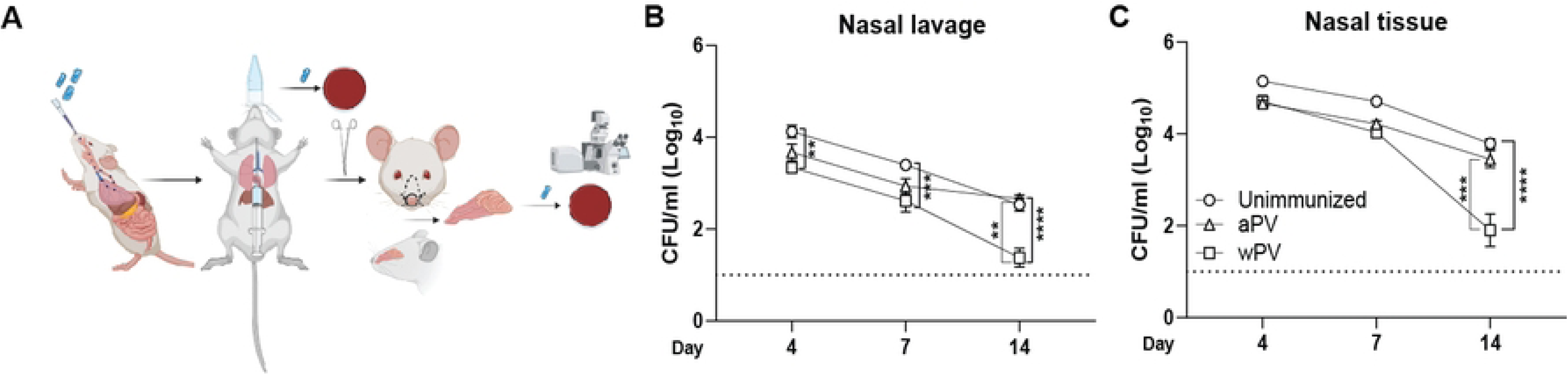
aPV immunization does not protect against *Bp* colonization and biofilm formation in the nasal cavity. (A) Diagram showing bacterial inoculation and tissue harvest. *Bp* CFUs in the (B) NL and (C) NT at days 4, 7, and 14 post challenge of unimmunized, aPV-immunized, and wPV-Cingiimmunized mice. Dotted line notes the limit of detection. Data shown are mean ± SEM (N=5-9) **P<0.01, ***P<0.001, ****P<0.0001.

We then determined the distribution of *Bp* in mice immunized with standard aPV and wPV. C57BL/6 mice were immunized intramuscularly (IM) on day 0 and day 28 with 1/10^th^ human dose of the aPV Boostrix^®^, or 1/10^th^ human dose of wPV, and challenged IN with Bp536 on day 42 (14 days post-boost). We quantified *Bp* burden in the NL and NT on days 4, 7, and 14 post-challenge. Bacterial load was similar in unimmunized and aPV immunized mice in the NL (**Fig. 1B**) and the NT (**Fig. 1C**) at all the time points evaluated. Over the entire infection time course, the bacterial load was significantly reduced in the NL of wPV immunized mice compared to unimmunized or aPV immunized mice (**Fig. 1B**). In contrast, bacterial burden was similar in the NT of all groups until day 14 post-challenge when wPV immunized mice had a significant reduction in bacterial burden compared to both unimmunized mice and aPV immunized mice (**Fig. 1B**). Thus, only wPV immunization reduces nasal bacterial burden although the nose is not completely cleared. Furthermore, quantification of bacteria in the NL and NT together is necessary since evaluation of the NL alone vastly underestimates the total burden.

We showed previously that *Bp* forms biofilms in the nasal cavity of naïve mice (11, 12). However, it is not known whether immunization with pertussis vaccines prevents the formation of biofilms. We collected the nasal septa from a separate group of naïve and immunized challenged mice and used confocal laser scanning electron microscopy (CLSM) to visualize the morphology of *Bp* remaining after NL collection. CLSM showed that *Bp* forms biofilms in unimmunized challenged mice as evident by multi-layered, large bacterial aggregates (green) adhered to the nasal epithelium (red) (**Fig. 2A and 2E**). Three-dimensional reconstructions of z-section image stacks created by IMARIS software revealed large irregularly shaped *Bp* clusters (**Fig. 2E**). Similar to unimmunized mice, *Bp* biofilms were also visualized in the NT of aPV immunized mice (**Fig. 2B and 2F**). In contrast, confocal microscopy of wPV immunized mice (**Fig. 2C and 2G**) resulted in barely detectable staining for bacterial cells, that was similar to naïve mice (**Fig. 2D**) and showed comparatively fewer and smaller bacterial aggregates than in unimmunized or aPV immunized challenged mice.

**Fig. 2.**
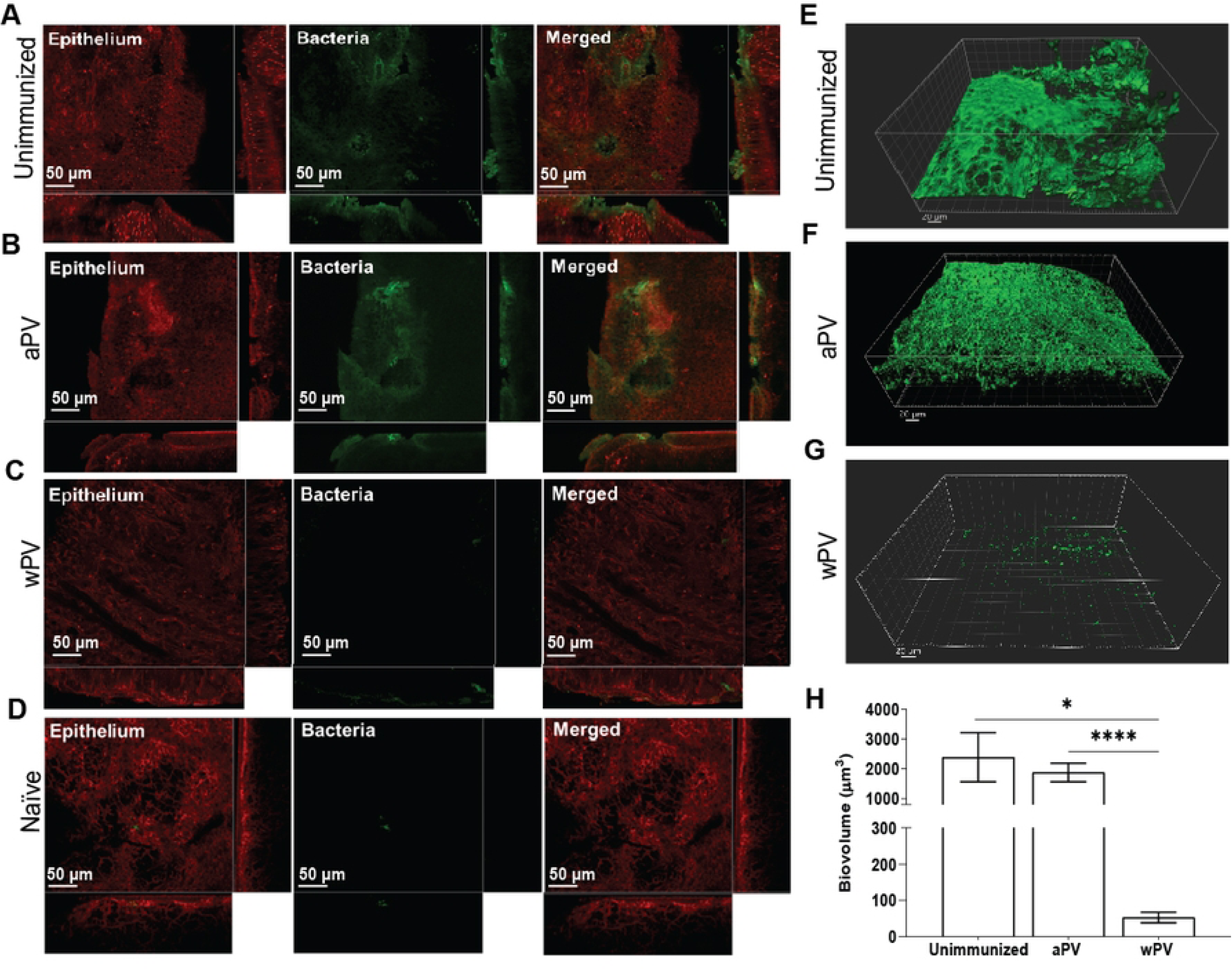
aPV immunization does not protect against *Bp* biofilm formation of the nasal cavity. CLSM analysis of tissues on day 7 post-challenge on nasal septa collected after lavage on images taken from 2-3 representative areas for each mouse (N=4-5) of (A) unimmunized challenged, (B) aPV immunized challenged, (C) wPv immunized challenged, and (D) naïve mice. IMARIS software was utilized to visualize *z*-stacks projections of (E) unimmunized, (F) aPV, and (G) wPV immunized challenged mice; *x-z, y-z,* and *xz* planes and *Z*-stack images were processed and analyzed by BiofilmQ software to determine the volume of the biofilms.(H) x-z, y-z, and xz planes and Z-stack images were processed and analyzed by BiofilmQ software to determine the biovolume of the biofilms. The bio-volume is defined as the number of biomass pixels in all images of a stack multiplied by the voxel size [(pixel size)x x (pixel size)y x (pixel size)z] and divided by the substratum area of the image stack. The resulting value is biomass volume divided by substratum area (µm3/µm2). Bio-volume represents the overall volume of the biofilm and provides an estimate of the biomass in the biofilm. Bars indicate mean ± SEM of at least one representative independent experiment with 4-5 mice. *P<0.05, ****P<0.0001 by t-test.

We quantified the biofilms biovolume using IMARIS software (**Fig. 2H**). Analysis of the NT after NL collection showed comparable bacterial biovolume between unimmunized (2,391.03±823.26 µm^3^), and aPV immunized mice (1,881.76±314.50 µm^3^) at 7 days post challenge (**Fig. 2H**). Strikingly, wPV mice showed 44.76-fold less biovolume in the nasal septum (53.41±14.55 µm^3^) compared to unimmunized and 35-fold less biovolume compared to aPV-immunized mice (**Fig. 2H**). Thus, *Bp* biofilm formation is drastically reduced in wPV immunized mice but is maintained in the noses of aPV immunized mice.

### Antibodies are primarily localized in the nasal tissues

To determine the nasal compartment where antibodies are primarily localized, we investigated the nasal serological responses in the NL and NT on day 14 post-challenge of unimmunized, aPV or wPV immunized mice. We compared responses elicited to the *Bp* protein FHA, which elicits antibodies in mice following aPV or wPV immunization (7) and to whole *Bp* bacteria. Notably, detectable antibody titers were found in the NT, but not in the NL (**Fig 3**). FHA-specific IgG was detected in all three groups (**Fig. 3**) and was higher in aPV and wPV immunized groups compared to unimmunized challenged mice. Anti-*Bp* specific IgG responses were also higher in immunized animals (**Fig. 3B**) and were localized predominantly within the NT. Anti-FHA specific IgA was detected in the NT in all groups but was not increased in aPV and wPV immunized mice compared to unimmunized challenged mice (**Fig. 3C**). However, *Bp*-specific IgA in the NT was elevated in wPV immunized animals compared to aPV immunized and unimmunized mice (**Fig. 3D**). Thus, most of the *Bp*-specific antibody responses are concentrated within the NT and not within the NL.

**Fig. 3.**
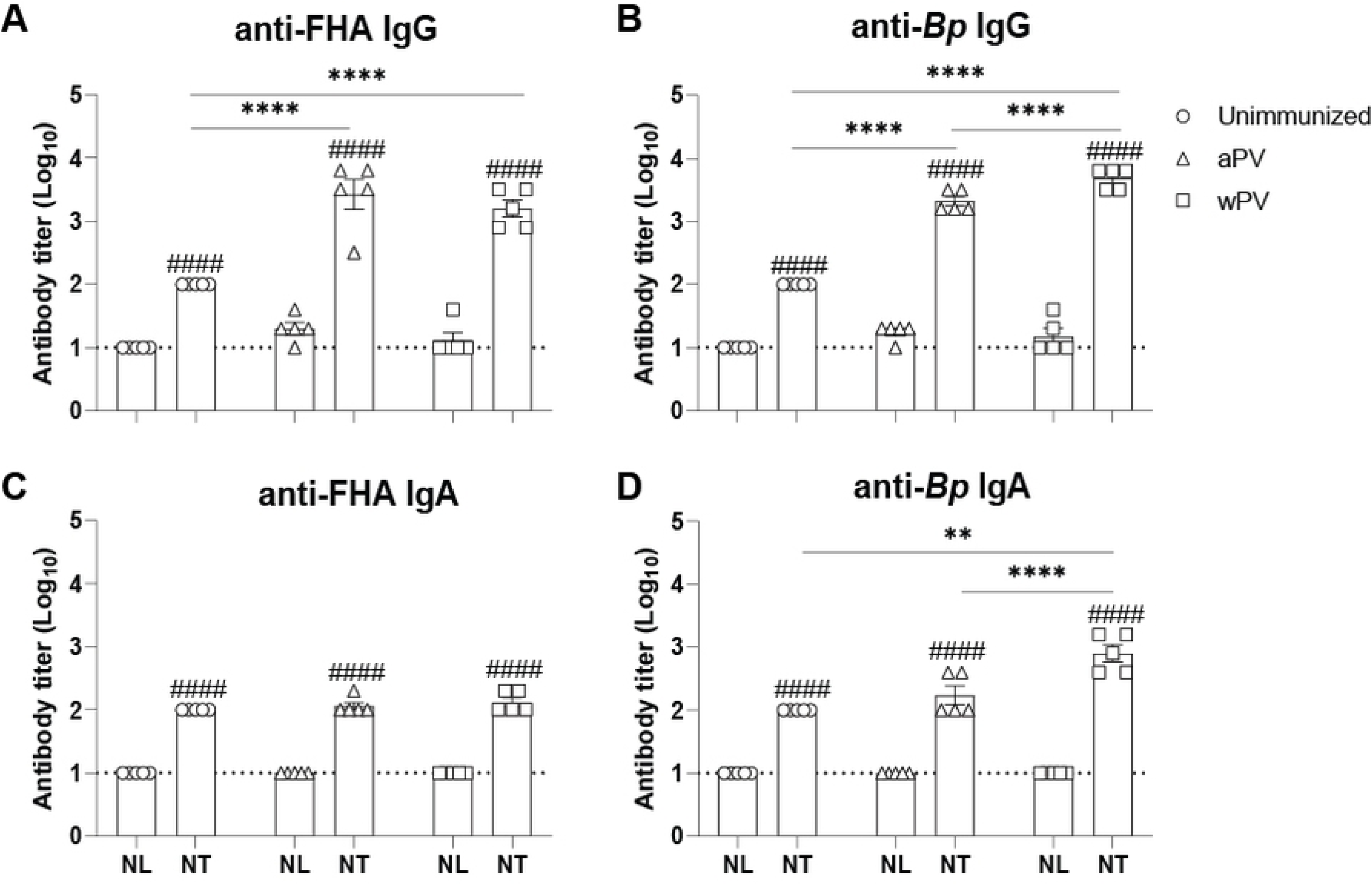
Nasal lavage underrepresents the antibody responses to immunization in the nasal cavity. ELISA for (A) anti-FHA specific and (B) anti-*Bp* specific IgG and (C) anti-FHA specific and (D) anti-Bp specific IgA antibodies in the nasal cavity (N=5-9). Mean ± SEM with the dotted line showing the limit of detection. ^####^, antibody responses in the NL compared to the NT. P<0.0001. **P<0.01, ****P<0.0001 between immunization groups.

### Immune cells are recruited to the nose following *Bp* challenge of wPV immunized mice

CD4+ T cells that produce IL-17 are critical for reduction of *Bp* bacterial burden from the nose (7). We used flow cytometry to enumerate the CD4+ T cells in the nose following aPV and wPV vaccination and challenge. Cell counts in the NL were too few to permit accurate enumeration of cell types. However, 100-fold more cells were recovered from the NT compared to the NL (**Fig. S2**). On day 7 post-challenge, CD4+ T cells in the NT of aPV immunized were similar to naïve and unimmunized challenged mice (**Fig. 4A**). wPV immunized mice had significantly higher numbers of CD4+ T cells in the NT (**Fig. 4A**) that was further amplified by day 14 post-challenge (**Fig. 4A**). At this time point, CD4+ T cells were also significantly increased in the NT of unimmunized *Bp* infected mice compared to naïve mice, demonstrating that infection alone recruits T cells to the mucosa.

**Fig. 4.**
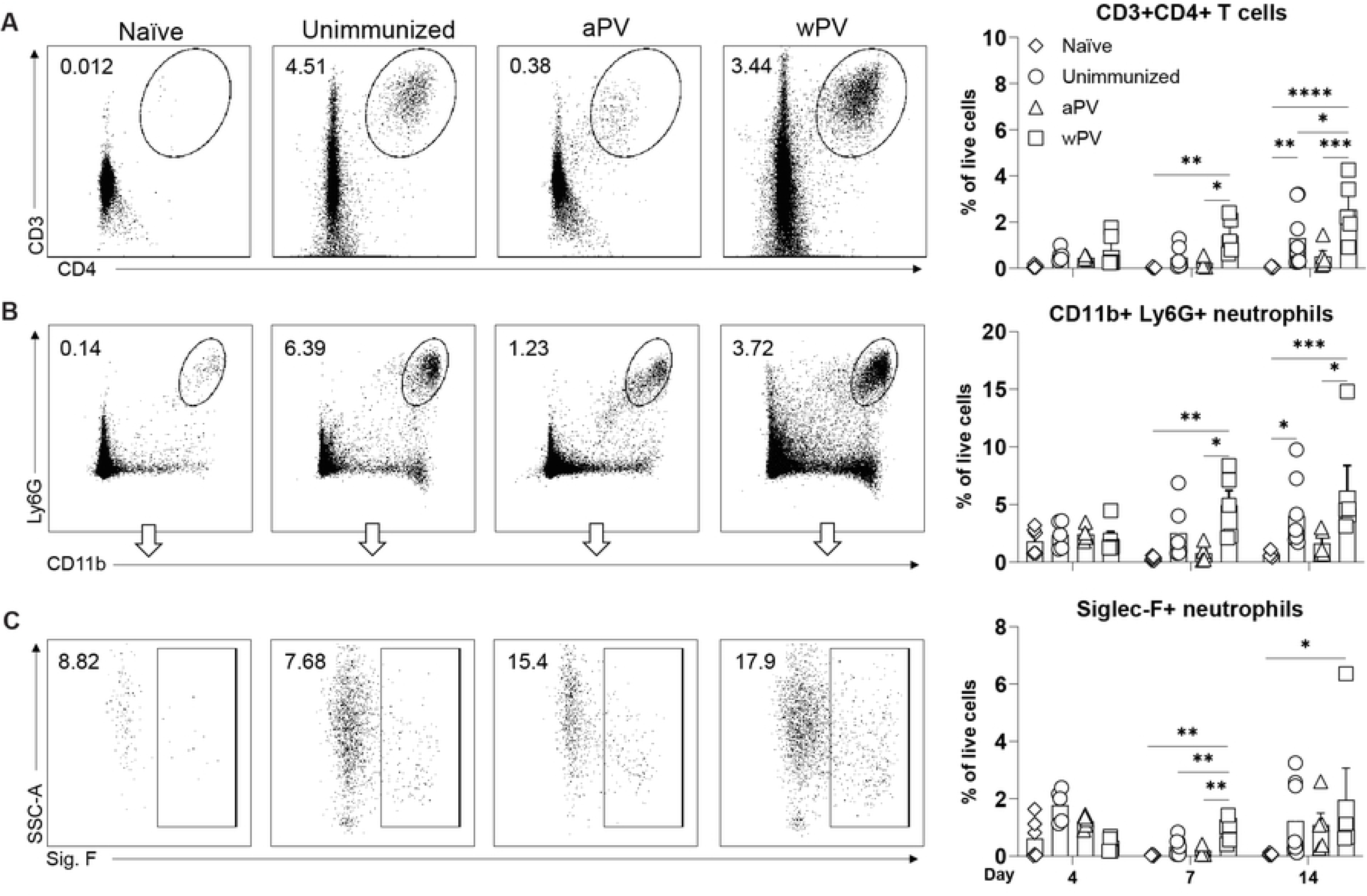
CD4+ T cells and Siglec-F+ neutrophils are recruited to the nasal cavity following *Bp* challenge of wPV immunized mice. Flow cytometry was used to determine the cellular response in the nasal cavity following immunization and challenge (N=5-6) at days 4, 7, and 14 post-challenge. (A) Representative gating and percentage of live cells that are CD3+, CD4+ T cells in the NT. Representative gating and percentage of live cells of (B) CD11b+Ly6G+ neutrophils and CD11b+Ly6G+Siglec-F+ neutrophils. Mean ± SEM are shown. *P<0.05, **P<0.01, ***P<0.001, ****P<0.0001.

In unimmunized mice, IL-17 producing T cells recruit neutrophils, particularly the Siglec-F+ subset to the infection site which clear *Bp* from the nasal cavity (17). On day 7 post-challenge, CD11b+ Ly6G+ neutrophils were significantly increased in wPV immunized mice compared to unimmunized and aPV immunized mice (**Fig. 4B**). Unimmunized mice displayed delayed kinetics with a significant increase of CD11b+ Ly6G+ neutrophils observed by day 14 post-challenge compared to naïve mice. (**Fig. 4B**). The CD11b+Ly6G+Siglec-F+ subset was significantly increased in wPV immunized mice at day 7 post challenge compared to unimmunized and aPV immunized mice (**Fig. 4C**) and remained elevated until day 14 post-challenge. These data show that Siglec-F+ neutrophils are key effectors that are recruited to the nose by wPV immunization but not by aPV immunization.

Macrophages are also recruited to the lungs of *Bp* challenged mice and mediate bacterial clearance (18, 19). Ly6C+ positive macrophages are recruited to the nose following aerosol challenge and have a proinflammatory phenotype, while Ly6C-macrophages maintain tissue homeostasis (20). We quantified the CD11b+ F4/80+ macrophages elicited by immunization and challenge. Total CD11b+ F4/80+ macrophages did not increase over the course of infection in any of the groups of challenged mice (**Fig. 5A**). In contrast, CD11b+ F4/80+ Ly6C+ macrophages were increased in the NT of wPV immunized mice compared to unimmunized and aPV-immunized mice at all time points (**Fig. 5B**). Thus, following IN *Bp* challenge, proinflammatory macrophages are recruited to the nose in wPV immunized mice, while this cell population does not increase in aPV immunized mice.

**Fig. 5.**
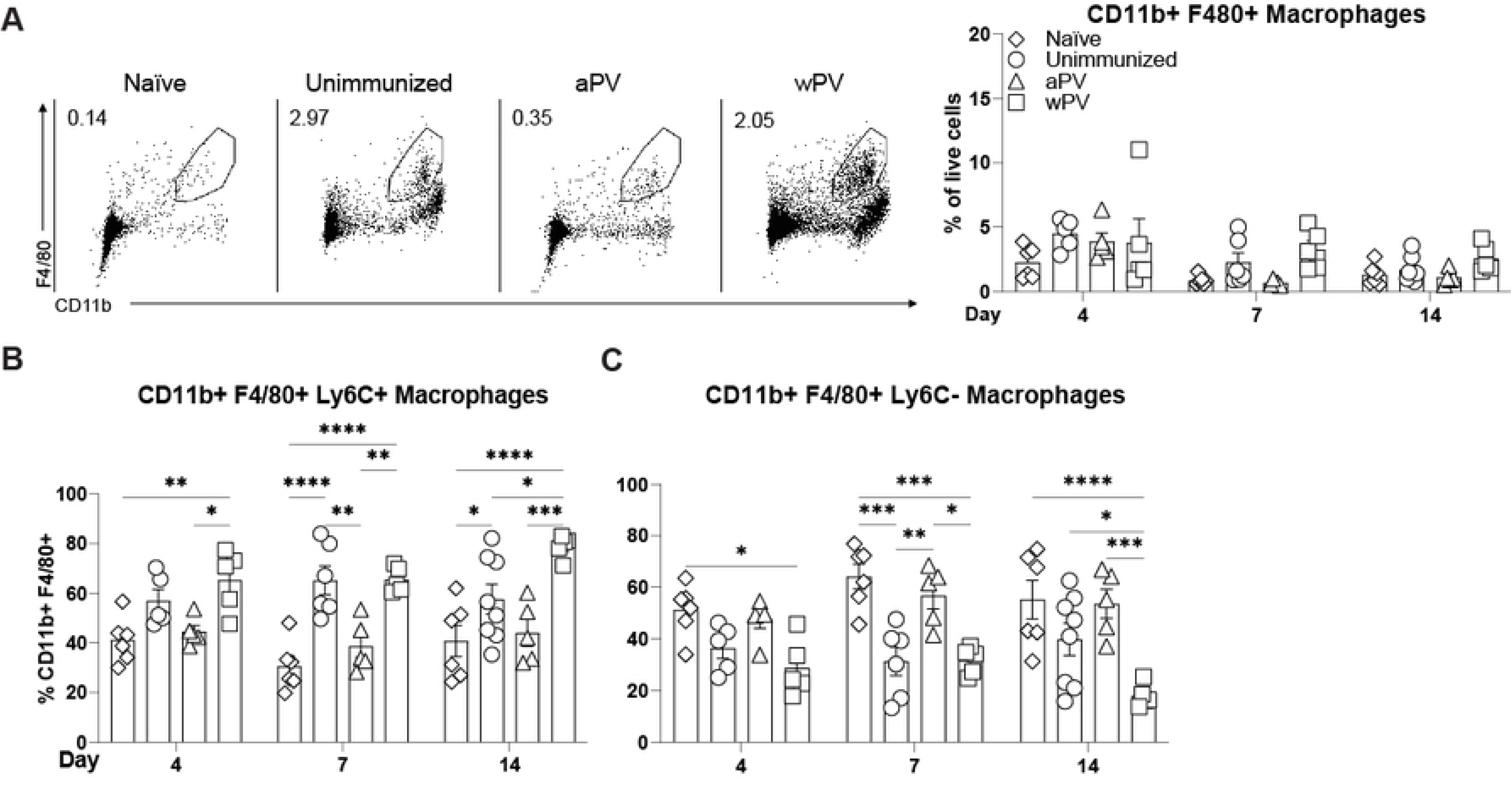
wPV immunization elicits Ly6C+ macrophages to the nasal cavity following *Bp* challenge. Flow cytometry was used to determine the CD11b+ F4/80+ macrophages in the NT following immunization and challenge (N=5-6) at days 4, 7, and 14 post-challenge. (A) Representative gating and percentage of live cells that are CD11b+ F4/80+ macrophages in the NT. Percentage of (B) CD11b+F4/80+Ly6C– and (C) CD11b+F4/80+Ly6C+ macrophages. Mean ± SEM. *P<0.05, **P<0.01, ***P<0.001, ****P<0.0001.

In contrast, the CD11b+ F4/80+ Ly6C-macrophages significantly increased in aPV immunized mice following *Bp* challenge compared to wPV immunized and *Bp* challenged mice (**Fig. 5C**), suggesting that aPV immunization elicits innate immune cells that prevent bacterial clearance.

### Neutrophil chemoattractants are produced in the NT of wPV immunized mice post-challenge

Chemokines are small molecules that direct specific leukocyte migration (21). Here, we evaluated CXCL1 and CXCL2 in the NT at day 14 post challenge by ELISA, as both chemokines regulate homing of neutrophils to the site of infection (22). Unimmunized and aPV immunized mice had minimally detectable amounts of CXCL1 in the NT, while wPV immunized mice had a significant increase in CXCL1 in the NT compared to naïve and aPV immunized mice (**Fig. 6A**). There was a slight but insignificant increase in in the NT of wPV immunized mice at day 14 post challenge (**Fig. 6B**). Thus, following *Bp* challenge, neutrophil chemoattractants are increased in the nasal cavity of wPV immunized but not aPV immunized mice.

**Fig. 6.**
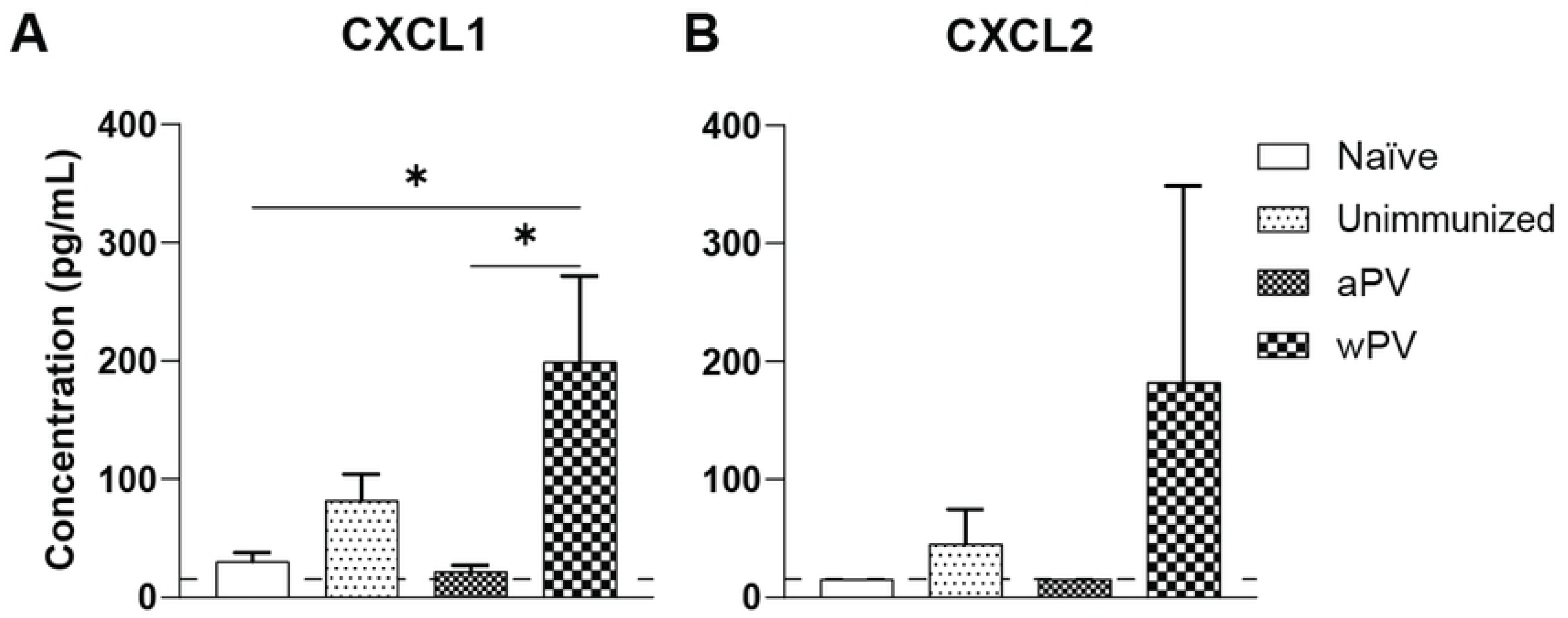
Chemokine CXCL1 was detected in the NT of wPV immunized mice following *Bp* challenge. ELISA was used to detect CXCL1 and CXCL2 in the NT of naïve, unimmunized, aPV and wPV immunized and challenged mice (N=5-9) at day 14 post challenge in the NT. Dotted line represents the limit of detection. Data is represented as mean ± SEM. *P<0.05.

## Discussion

While it is well recognized that aPV fail to prevent *Bp* nasal colonization, the mechanism responsible for maintenance of the nasal reservoir is not known. Here, we provide insights into this long-standing observation. CLSM showed that *Bp* resides in the nasal septum as biofilms in naïve and aPV immunized mice by day 7 post-challenge. In contrast, wPV immunization prevented biofilm formation. *Bp* biofilm formation is a key pathogenic mechanism (23) that facilitates *Bp* survival and persistence in the respiratory tract in mice (24, 25). These structures are like those observed on human respiratory epithelium (26). We propose that maintenance of *Bp* as biofilms in the nose results in aPV immunized populations that are capable of asymptomatic transmission of *Bp* to unvaccinated or partially vaccinated infants or immune compromised populations. Since aPV immunization does not reduce bacterial burden in the nose, bacterial biofilm formation is not prevented, which permits *Bp* persistence. Our data are consistent with previous findings (5, 7) that the nasal cavity of aPV immunized mice remains colonized. These studies were conducted in Balb/c mice which generate T_H_2 polarized immune responses. Our studies in T_H_1 polarized C57BL/6 mice yielded the same results, confirming that the bacterial maintenance in aPV immunized animals is independent of genetic background and inherent immune polarization.

Our comparison of NL and NT shows that NL alone vastly underrepresents the *Bp* burden in the nasal cavity during infection. Quantification of *Bp* CFUs at various time points post-challenge of unimmunized, aPV-immunized, and wPV-immunized mice showed that most of the bacteria are localized in the NT, and enumeration solely in the lavage represented less than 90% of the bacterial load. Importantly, bacterial load in the NT was two logs higher than in the NL. wPV immunization elicited better reduction of *Bp* yet not eliminating *Bp* from the nasal tissue entirely. NL collects bacteria weakly associated to the epithelium or freely floating in the nasal cavity. However, nasal cavity tissue extraction is necessary for comprehensive evaluation of bacterial load (27, 28). The nasal cavity consists of nasal turbinates, NALT, nasal septum, and ciliated epithelial cells all capable of trapping bacteria and facilitating bacterial colonization (29–31). Nasal turbinates regulate the temperature of the inspired air and filter out foreign substances, while mucosal immune responses are initiated in the NALT (32). Fluorescence microscopy confirmed the CFU enumeration results and showed that *Bp* adheres to the septum and NT (25, 26), with the potential to form biofilms.

Serological responses to *Bp* are used to determine vaccine efficacy and protection in humans (33)and animals (13, 34)and are quantified in nasal secretions or nasal lavage. We found that antibodies in the NL were at the limit of detection, with antibodies in the NT significantly (∼1-2 log) increased in wPV immunized mice. aPV and wPV immunization elicited FHA-specific IgG antibodies that accumulated in NT. At day 14 post-challenge, anti-*Bp* specific IgG antibodies were detected in the NT of all groups of mice. While mucosal immunization elicits IgA responses in the nose (13, 35, 36), *Bp*-specific IgA antibodies in the NT were only detected in wPV immunized mice, suggesting that the systemic immune response elicited by wPV is recruited to the mucosa upon bacterial challenge. Thus, accurate quantification of antibodies in the respiratory tract necessitates evaluation of the NT in addition to the NL.

CD4+ T cells were increased in the NT following bacterial challenge of wPV immunized mice, while challenged aPV immunized mice did not have significant numbers of CD4+ T cells in the NT. aPV immunization elicits T_H_2 polarized systemic T cells specific for the aPV antigens, FHA, Prn and PT (15, 37–41). However, these may not be the immunodominant *Bp* antigens presented following infection [Shamseldin, Hall, Hernandez et al, submitted], and thus the aPV antigen specific T cells may not be mobilized to respond to the infection (42, 43).

The neutrophil chemoattractant CXCL1 was significantly increased only in the NT of wPV immunized mice and correlated with recruitment of Siglec-F+ neutrophils to the nose of wPV immunized mice. Total neutrophils and the Siglec-F+ fraction did not increase in aPV immunized mice and correlated with maintenance of *Bp* in the NT. Interestingly, the Ly6C+ inflammatory macrophage subset was also increased in the NT of wPV immunized mice, suggesting a potential role for this cell population in elimination of *Bp* from the upper respiratory tract. We are investigating the relative importance of these two phagocytes in another project. Thus, accurate evaluation of bacterial burden and the immune response generated by vaccination necessitates evaluation of the nasal tissues where both bacteria and humoral and cellular components become sequestered.

aPV and wPV are both administered IM and elicit serum antibodies and systemic T cell responses. How immune cells are recruited to the mucosa following *Bp* challenge of wPV immunized animals is an important, open question that warrants further investigation. While the administration route is the same, the vaccine formulations are distinct. wPV is made by chemically inactivating *Bp*, and contains many potential antigens, while aPV comprise only 3-5 antigens, including pertussis toxin (PT), filamentous hemagglutinin (FHA), pertactin (Prn), and fimbriae 2/3 (fim2/3) adjuvanted with alum, which limits the diversity of the immune response. Natural infection and wPV immunization (44, 45)elicit long-lived T_H_1/T_H_17 responses while aPV elicit relatively short-livedT_H_2-polarized immunity (46–48). The waning of aPV immune responses (49–51)is correlated with the emergence of vaccine escape strains globally (52). These produce hyper biofilms *in vitro* (24) and may have increased capacity to form biofilms in the respiratory tract.

Together, our data show that aPV and wPV immunization has distinct consequences on bacterial clearance and biofilm formation in the upper respiratory tract. These results further suggest that next generation vaccine formulations should elicit sustained T_H_1/T_H_17 polarized cellular mucosal immunity. Several studies including our own (15, 53) show that subunit vaccines that contain T_H_1/T_H_17 polarizing adjuvants, where at least one dose is delivered IN elicit this immune phenotype. Consequently, bacterial burden in the nose is significantly reduced. Whether these formulations prevent *Bp* aggregation, biofilm formation and subsequent transmission is an important, unanswered question. These vaccines also do not include antigens that promote nasal colonization. Only one such factor, the polysaccharide Bps (11) has been identified that is critical for biofilm formation and nasal colonization of immunocompetent mice. Bps is not a component of current subunit acellular pertussis vaccines. Thus, novel *Bp* vaccines which incorporate adjuvants that elicit T_H_17 polarized immunity and antigens involved in nasal cavity colonization will prevent biofilm formation and thereby control *Bp* persistence and transmission. This work also provides a framework to evaluate the efficacy of vaccines against other bacterial and viral pathogens where nasal colonization regulates transmission.

## Materials and Methods

### *Bp* growth conditions

*Bordetella pertussis* strain Bp536 (54) was grown on Bordet Gengou (BG) plates (Difco, (Ref. 248200) containing 10% defibrinated sheep’s blood (Hemostat) and 100µg/ml streptomycin for 4 days at 37°C. Following incubation, *Bp* was transferred to Stainer-Scholte medium supplemented with 1mg/ml (2,3,6-tri-O-methyl)-β-cyclodextrin Heptakis (Sigma-Aldrich, Cat. H0513-5G) and incubated in a rolling drum at 180 rpm at 37°C for 24 hours. When the culture reached ∼OD_600_=1 the bacteria was diluted in sterile, endotoxin free 1x PBS for intranasal inoculation.

### Immunization

The acellular pertussis vaccine, Boostrix (GSK) and wPV (DTP) (NIBSC) were diluted to 1/10^th^ human dose in 50µl of endotoxin free 1X PBS/dose. C57BL/6 mice (6-12 weeks old) were intramuscularly (IM) immunized on day 0 and boosted on day 28 in the forelimb with the same dose.

### *Bp* challenge

Mice were anesthetized with 2.5% isoflurane/O2 for bacterial inoculation. Immunized mice and unimmunized age matched controls were inoculated IN with *Bp* diluted to 5×10^5^ CFU/mouse in 50µl, divided equally between both nares.

### Tissue collection, *Bp* enumeration, and flow cytometry

Following euthanasia, the nares were flushed with 1 mL PBS using a syringe attached to a 20G X 1.25 catheter (Exel International Ref. 26742). The solution was injected into the bottom of the nasal cavity above the trachea and collected from the tip of the nose. Then, nasal tissues were excised by cutting 1-2mm of the tip of the nose. Sagittal cuts were made on each side of the nasal septum exposing the nasal associated tissue. The nasal septum, NALT, and turbinates were harvested and were enzymatically digested with GentleMACS lung dissociation kit (Miltenyi) according to the manufacturer’s instructions. Before filtration, NL and NT homogenates were diluted and plated on BG plates to enumerate CFUs. The NT cell suspensions were filtered through a 40-μm cell strainer (Fisherbrand Cat. 22363547), and supernatant was collected and stored at – 80°C for serological and chemokine analysis. Red blood cells were lysed with ACK lysis buffer (Gibco, Ref. A10492-01). The NL and NT cell populations were counted by trypan blue exclusion using a hemacytometer to determine the number of cells collected in each fraction. NT cells were incubated with LIVE/DEAD Aqua (Invitrogen, Cat. L34966), followed by incubation with α-CD16/CD32 FcγRIII (Thermo Fisher Scientific, Cat. 14-0161-86) to block nonspecific binding. Antibodies used to identify specific cell populations are listed in Table 1. Pooled samples from mice in each group were used as unstained negative controls and were stained with single antibodies as fluorescence minus one (FMO) samples for compensation. Samples were acquired on a Cytek Aurora spectral flow cytometer and data were analyzed with FlowJo software.

**Table 1.** Flow cytometry antibody panel for evaluation of immune cell populations of the nose.

### Confocal Laser Scanning Microscopy and image analysis

At 7 days post-challenge, mice were euthanized, and the nasal septum and trachea were harvested after nasal lavage. The tissues were fixed overnight at 4 °C in 4% methanol-free paraformaldehyde (ThermoFisher Scientific, cat no. U01H501), and then washed three times with 1xPBS for 30 min. To permeabilize the epithelium, organs were incubated with 0.1% Triton X-100 (Fisher Bioreagents, cat no. BP151-100) for 15 min at room temperature (RT). Samples were then blocked with 5% normal goat serum (Abcam, cat no. ab7481) in 1% BSA for 30 min at RT, and then washed once for 30 min with 1X PBS. To block endogenous IgG, samples were treated with unconjugated affinity purified F(ab) fragment IgG anti-mouse (H+L) for 1 hr at RT (0.1 mg/ml, Abcam, cat no. ab6668). Mouse epithelium was stained with Alexa 633-conjugated phalloidin (Invitrogen, cat no. a22284) per manufacturer’s instructions. To label *Bp*, samples were incubated overnight at 4 °C with immune serum from mice immunized with aPV (1:1,000), then washed 6X for 1 hr each, and incubated for 2 hr at RT with goat IgG anti-mouse secondary antibody conjugated to Oregon 488 (1 µg/ml, Invitrogen cat no. O-6380). Tissues were mounted on 1 mm thick slides using Prolong Gold antifade (Invitrogen, cat no. P36934) and 22×22 mm cover glasses. Z-stack images were acquired by Confocal Scanning Electron Microscopy (CLSM) at 0.5-µm z-intervals using an Olympus FV300 confocal microscope. IMARIS software (Biplane) was used to visualize z-stacks projections. For this, gray values were adjusted to 1.30 gamma correction. Z-stacks were then processed and analyzed by BiofilmQ software (MATLAD vR2019b) (55) to determine the biovolume of the biofilms by using automatic Otsu’s thresholding. Five samples were analyzed per group.

### ELISA analysis

Enzyme-linked immunosorbent assay (ELISA) was used to quantify antigen-specific antibody titers and chemokines in nasal lavage and nasal tissue supernatant. Protein high-binding 96 well plates (Corning #9018) were coated with 1µg/ml purified FHA (produced by Dr. Jennifer Maynard, UT-Austin). To determine antibody responses to the whole bacteria, the plates were coated with intact *Bp* bacteria o/n at 4°C and then washed with 0.5% PBS Tween 20. Samples were blocked with ELISA blocking diluent (Invitrogen) for 2 hr at 37°C then washed 3X. Nasal lavage and nasal tissue samples were diluted and added to the antigen coated plates and incubated o/n at 4°C. Plates were washed and probed with either anti-mouse IgG (Southern Biotech, Cat. 2040-05) or anti-mouse IgA (Invitrogen Cat. 292318) HRP for 1 hr at RT. Plates were washed and developed with TMB (BioLegend, Cat. 421501). The reaction was stopped with 2N H_2_SO_4_ and plates were read at an AB_450_ nm. CXCL1 (R&D ref. DY453) and CXCL2 (R&D ref. DY452) were determined in the nasal tissue supernatant by per manufacturer’s protocol. Briefly, 96-well microplates were coated with diluted capture antibody o/n at RT, then blocked with reagent diluent for 1hr at RT. The plate was washed, and NT supernatant was added and incubated at RT for 2hrs. The chemokines were detected using biotinylated anti-mouse secondary antibody and streptavidin-HRP. Plates were developed using TMB and the reaction was stopped with 2N H_2_SO_4_. Plates were read at AB_450_ nm.

## Statistical analysis

GraphPad Prism 9 software was used to generate graphs and analyze data. Unpaired student’s t-test was used to compare bacterial burden in the nasal lavage and nasal tissue on the same day between 2 groups, and for biofilm biovolume comparisons. One-way and two-way analysis of variance (ANOVA) with multiple comparisons and Tukey’s *post hoc* test was used for statistical comparison of data between immunized and unimmunized *Bp* challenged mice at each time point.

## Ethics statement

All animal work was approved by Institutional Animal Care and Use Committee (IACUC) at The Ohio State University (OSU), protocol number: 2017A00000090-2.

## Data availability

All raw data will be made available upon request.

## Acknowledgments

We thank The Ohio State University Campus Microscopy Imaging Facility (CMIF) for use of the confocal laser scanning microscope, and for the use of the IMARIS software. We also thank Dr. Jennifer Maynard for producing the purified FHA protein.

## Figure Legends

**Fig. S1.**
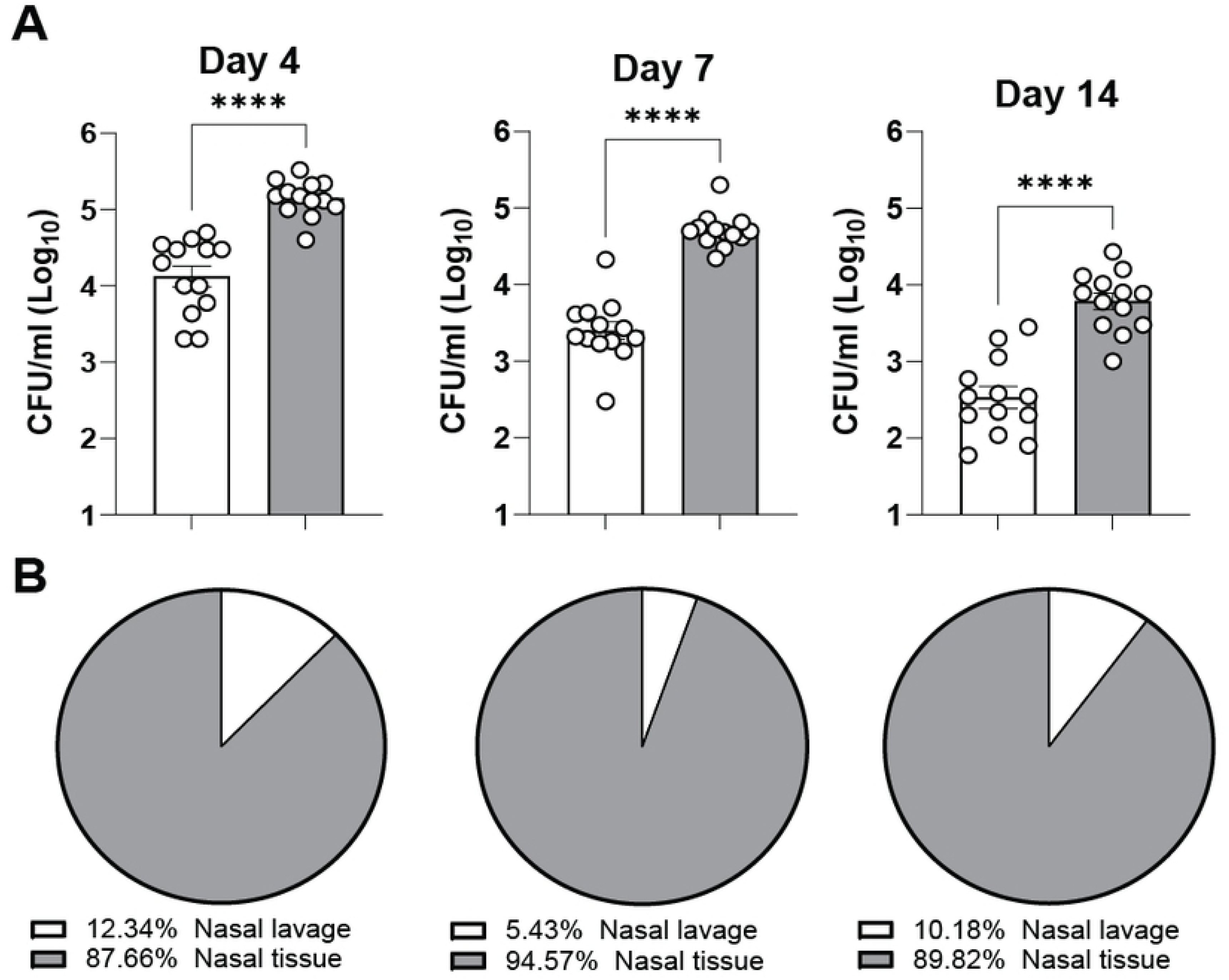
Nasal lavage alone underrepresents *Bp* burden in the nasal cavity. (A) *Bp* CFUs in the nasal lavage and nasal septum of unimmunized mice challenged with *Bp* and harvested at days 4, 7, and 14 post challenge. Data shown as mean ± SEM of 3 different experiments (N=13). CFUs in nasal lavage compared to the nasal associated tissue were determined by an unpaired t-test. ****P<0.0001. (B) Percentages of total bacteria that was collected in the NL compared to the bacterial burden in the NT.

**Fig. S2.**
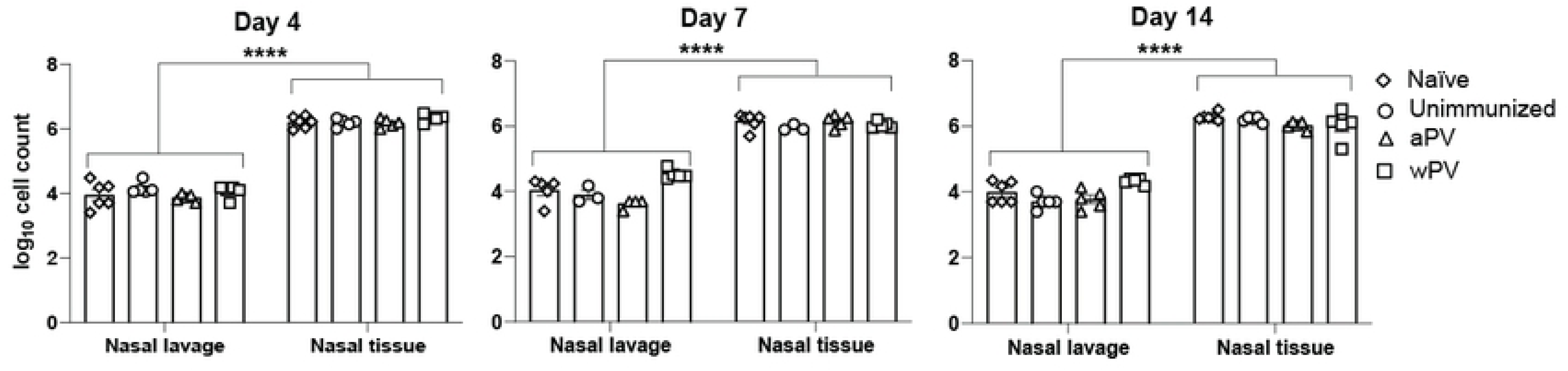
Nasal lavage largely underrepresents the cellular responses in the nasal cavity. Following nasal lavage and nasal tissue collection and processing, total cell counts were determined for flow cytometry. Mean ± SEM (N= 5-6). ****P<0.0001.

## References

1. Feldstein LR, Mariat S, Gacic-Dobo M, Diallo MS, Conklin LM, Wallace AS. Global Routine Vaccination Coverage, 2016. MMWR Morb Mortal Wkly Rep. 2017;66(45):1252–5.

2. Cody CL, Baraff LJ, Cherry JD, Marcy SM, Manclark CR. Nature and rates of adverse reactions associated with DTP and DT immunizations in infants and children. Pediatrics. 1981;68(5):650–60.

3. Sato Y, Sato H. Development of acellular pertussis vaccines. Biologicals. 1999;27(2):61–9.

4. Esposito S, Stefanelli P, Fry NK, Fedele G, He Q, Paterson P, et al. Pertussis Prevention: Reasons for Resurgence, and Differences in the Current Acellular Pertussis Vaccines. Front Immunol. 2019;10:1344.

5. Holubova J, Stanek O, Brazdilova L, Masin J, Bumba L, Gorringe AR, et al. Acellular Pertussis Vaccine Inhibits Bordetella pertussis Clearance from the Nasal Mucosa of Mice. Vaccines (Basel). 2020;8(4).

6. Warfel JM, Zimmerman LI, Merkel TJ. Acellular pertussis vaccines protect against disease but fail to prevent infection and transmission in a nonhuman primate model. Proc Natl Acad Sci U S A. 2014;111(2):787–92.

7. Wilk MM, Borkner L, Misiak A, Curham L, Allen AC, Mills KHG. Immunization with whole cell but not acellular pertussis vaccines primes CD4 T(RM) cells that sustain protective immunity against nasal colonization with Bordetella pertussis. Emerg Microbes Infect. 2019;8(1):169–85.

8. Althouse BM, Scarpino SV. Asymptomatic transmission and the resurgence of Bordetella pertussis. BMC Med. 2015;13:146.

9. de Graaf H, Ibrahim M, Hill AR, Gbesemete D, Vaughan AT, Gorringe A, et al. Controlled Human Infection With Bordetella pertussis Induces Asymptomatic, Immunizing Colonization. Clin Infect Dis. 2020;71(2):403–11.

10. Zhang Q, Yin Z, Li Y, Luo H, Shao Z, Gao Y, et al. Prevalence of asymptomatic Bordetella pertussis and Bordetella parapertussis infections among school children in China as determined by pooled real-time PCR: a cross-sectional study. Scand J Infect Dis. 2014;46(4):280–7.

11. Conover MS, Sloan GP, Love CF, Sukumar N, Deora R. The Bps polysaccharide of Bordetella pertussis promotes colonization and biofilm formation in the nose by functioning as an adhesin. Mol Microbiol. 2010;77(6):1439–55.

12. Serra DO, Conover MS, Arnal L, Sloan GP, Rodriguez ME, Yantorno OM, et al. FHA-mediated cell-substrate and cell-cell adhesions are critical for Bordetella pertussis biofilm formation on abiotic surfaces and in the mouse nose and the trachea. PLoS One. 2011;6(12):e28811.

13. Boehm DT, Wolf MA, Hall JM, Wong TY, Sen-Kilic E, Basinger HD, et al. Intranasal acellular pertussis vaccine provides mucosal immunity and protects mice from Bordetella pertussis. NPJ Vaccines. 2019;4:40.

14. Raeven RHM, Rockx-Brouwer D, Kanojia G, van der Maas L, Bindels THE, Ten Have R, et al. Intranasal immunization with outer membrane vesicle pertussis vaccine confers broad protection through mucosal IgA and Th17 responses. Sci Rep. 2020;10(1):7396.

15. Yount KS, Hall JM, Caution K, Shamseldin MM, Guo M, Marion K, et al. Systemic priming and intranasal booster with a BcfA-adjuvanted acellular pertussis vaccine generates CD4+ IL-17+ nasal tissue resident T cells and reduces B. pertussis nasal colonization. Front Immunol. 2023;14:1181876.

16. Sato Y, Arai H. Leucocytosis-promoting factor of Bordetella pertussis. I. Purification and characterization. Infect Immun. 1972;6(6):899–904.

17. Borkner L, Curham LM, Wilk MM, Moran B, Mills KHG. IL-17 mediates protective immunity against nasal infection with Bordetella pertussis by mobilizing neutrophils, especially Siglec-F(+) neutrophils. Mucosal Immunol. 2021;14(5):1183–202.

18. Carbonetti NH, Artamonova GV, Van Rooijen N, Ayala VI. Pertussis toxin targets airway macrophages to promote Bordetella pertussis infection of the respiratory tract. Infect Immun. 2007;75(4):1713–20.

19. McGuirk P, Mahon BP, Griffin F, Mills KH. Compartmentalization of T cell responses following respiratory infection with Bordetella pertussis: hyporesponsiveness of lung T cells is associated with modulated expression of the co-stimulatory molecule CD28. Eur J Immunol. 1998;28(1):153–63.

20. Li YH, Zhang Y, Pan G, Xiang LX, Luo DC, Shao JZ. Occurrences and Functions of Ly6C(hi) and Ly6C(lo) Macrophages in Health and Disease. Front Immunol. 2022;13:901672.

21. Hughes CE, Nibbs RJB. A guide to chemokines and their receptors. FEBS J. 2018;285(16):2944–71.

22. Zhou C, Gao Y, Ding P, Wu T, Ji G. The role of CXCL family members in different diseases. Cell Death Discov. 2023;9(1):212.

23. Soane MC, Jackson A, Maskell D, Allen A, Keig P, Dewar A, et al. Interaction of Bordetella pertussis with human respiratory mucosa in vitro. Respir Med. 2000;94(8):791–9.

24. Cattelan N, Jennings-Gee J, Dubey P, Yantorno OM, Deora R. Hyperbiofilm Formation by Bordetella pertussis Strains Correlates with Enhanced Virulence Traits. Infect Immun. 2017;85(12).

25. Sloan GP, Love CF, Sukumar N, Mishra M, Deora R. The Bordetella Bps polysaccharide is critical for biofilm development in the mouse respiratory tract. J Bacteriol. 2007;189(22):8270–6.

26. Fullen AR, Gutierrez-Ferman JL, Rayner RE, Kim SH, Chen P, Dubey P, et al. Architecture and matrix assembly determinants of Bordetella pertussis biofilms on primary human airway epithelium. PLoS Pathog. 2023;19(2):e1011193.

27. Dubois V, Chatagnon J, Thiriard A, Bauderlique-Le Roy H, Debrie AS, Coutte L, et al. Suppression of mucosal Th17 memory responses by acellular pertussis vaccines enhances nasal Bordetella pertussis carriage. NPJ Vaccines. 2021;6(1):6.

28. Schenck LP, McGrath JJC, Lamarche D, Stampfli MR, Bowdish DME, Surette MG. Nasal Tissue Extraction Is Essential for Characterization of the Murine Upper Respiratory Tract Microbiota. mSphere. 2020;5(6).

29. Cingi C, Bayar Muluk N, Mitsias DI, Papadopoulos NG, Klimek L, Laulajainen-Hongisto A, et al. The Nose as a Route for Therapy: Part 1. Pharmacotherapy. Front Allergy. 2021;2:638136.

30. Davis JD, Wypych TP. Cellular and functional heterogeneity of the airway epithelium. Mucosal Immunol. 2021;14(5):978–90.

31. Sahin-Yilmaz A, Naclerio RM. Anatomy and physiology of the upper airway. Proc Am Thorac Soc. 2011;8(1):31–9.

32. Alvites RD, Caseiro AR, Pedrosa SS, Branquinho ME, Varejao ASP, Mauricio AC. The Nasal Cavity of the Rat and Mouse-Source of Mesenchymal Stem Cells for Treatment of Peripheral Nerve Injury. Anat Rec (Hoboken). 2018;301(10):1678–89.

33. Goodman YE, Wort AJ, Jackson FL. Enzyme-linked immunosorbent assay for detection of pertussis immunoglobulin A in nasopharyngeal secretions as an indicator of recent infection. J Clin Microbiol. 1981;13(2):286–92.

34. Jiang W, Wang X, Su Y, Cai L, Li J, Liang J, et al. Intranasal Immunization With a c-di-GMP-Adjuvanted Acellular Pertussis Vaccine Provides Superior Immunity Against Bordetella pertussis in a Mouse Model. Front Immunol. 2022;13:878832.

35. Solans L, Debrie AS, Borkner L, Aguilo N, Thiriard A, Coutte L, et al. IL-17-dependent SIgA-mediated protection against nasal Bordetella pertussis infection by live attenuated BPZE1 vaccine. Mucosal Immunol. 2018;11(6):1753–62.

36. Wolf MA, Boehm DT, DeJong MA, Wong TY, Sen-Kilic E, Hall JM, et al. Intranasal Immunization with Acellular Pertussis Vaccines Results in Long-Term Immunity to Bordetella pertussis in Mice. Infect Immun. 2021;89(3).

37. Ausiello CM, Urbani F, la Sala A, Lande R, Cassone A. Vaccine– and antigen-dependent type 1 and type 2 cytokine induction after primary vaccination of infants with whole-cell or acellular pertussis vaccines. Infect Immun. 1997;65(6):2168–74.

38. Jennings-Gee J, Quataert S, Ganguly T, D’Agostino R, Jr., Deora R, Dubey P. The Adjuvant Bordetella Colonization Factor A Attenuates Alum-Induced Th2 Responses and Enhances Bordetella pertussis Clearance from Mouse Lungs. Infect Immun. 2018;86(6).

39. Redhead K, Watkins J, Barnard A, Mills KH. Effective immunization against Bordetella pertussis respiratory infection in mice is dependent on induction of cell-mediated immunity. Infect Immun. 1993;61(8):3190–8.

40. Rowe J, Macaubas C, Monger TM, Holt BJ, Harvey J, Poolman JT, et al. Antigen-specific responses to diphtheria-tetanus-acellular pertussis vaccine in human infants are initially Th2 polarized. Infect Immun. 2000;68(7):3873–7.

41. Rowe J, Yerkovich ST, Richmond P, Suriyaarachchi D, Fisher E, Feddema L, et al. Th2-associated local reactions to the acellular diphtheria-tetanus-pertussis vaccine in 4– to 6– year-old children. Infect Immun. 2005;73(12):8130–5.

42. Ghani S, Feuerer M, Doebis C, Lauer U, Loddenkemper C, Huehn J, et al. T cells as pioneers: antigen-specific T cells condition inflamed sites for high-rate antigen-non-specific effector cell recruitment. Immunology. 2009;128(1 Suppl):e870–80.

43. Woodland DL, Kohlmeier JE. Migration, maintenance and recall of memory T cells in peripheral tissues. Nat Rev Immunol. 2009;9(3):153–61.

44. Lambert HJ. Epidemiology of a Small Pertussis Outbreak in Kent County, Michigan. Public Health Rep (1896). 1965;80(4):365–9.

45. Wendelboe AM, Van Rie A, Salmaso S, Englund JA. Duration of immunity against pertussis after natural infection or vaccination. Pediatr Infect Dis J. 2005;24(5 Suppl):S58–61.

46. Lugauer S, Heininger U, Cherry JD, Stehr K. Long-term clinical effectiveness of an acellular pertussis component vaccine and a whole cell pertussis component vaccine. Eur J Pediatr. 2002;161(3):142–6.

47. Simondon F, Preziosi MP, Yam A, Kane CT, Chabirand L, Iteman I, et al. A randomized double-blind trial comparing a two-component acellular to a whole-cell pertussis vaccine in Senegal. Vaccine. 1997;15(15):1606–12.

48. Tindberg Y, Blennow M, Granstrom M. A ten year follow-up after immunization with a two component acellular pertussis vaccine. Pediatr Infect Dis J. 1999;18(4):361–5.

49. Klein NP, Bartlett J, Fireman B, Baxter R. Waning Tdap Effectiveness in Adolescents. Pediatrics. 2016;137(3):e20153326.

50. Klein NP, Bartlett J, Rowhani-Rahbar A, Fireman B, Baxter R. Waning protection after fifth dose of acellular pertussis vaccine in children. N Engl J Med. 2012;367(11):1012–9.

51. Sheridan SL, Frith K, Snelling TL, Grimwood K, McIntyre PB, Lambert SB. Waning vaccine immunity in teenagers primed with whole cell and acellular pertussis vaccine: recent epidemiology. Expert Rev Vaccines. 2014;13(9):1081–106.

52. Jayasundara D, Lee E, Octavia S, Lan R, Tanaka MM, Wood JG. Emergence of pertactin-deficient pertussis strains in Australia can be explained by models of vaccine escape. Epidemics. 2020;31:100388.

53. Shamseldin MM, Kenney A, Zani A, Evans JP, Zeng C, Read KA, et al. Prime-Pull Immunization of Mice with a BcfA-Adjuvanted Vaccine Elicits Sustained Mucosal Immunity That Prevents SARS-CoV-2 Infection and Pathology. J Immunol. 2023;210(9):1257–71.

54. Relman DA, Domenighini M, Tuomanen E, Rappuoli R, Falkow S. Filamentous hemagglutinin of Bordetella pertussis: nucleotide sequence and crucial role in adherence. Proc Natl Acad Sci U S A. 1989;86(8):2637–41.

55. Hartmann R, Jeckel H, Jelli E, Singh PK, Vaidya S, Bayer M, et al. Quantitative image analysis of microbial communities with BiofilmQ. Nat Microbiol. 2021;6(2):151–6.

